# cfMIND: A read-level methylation framework for accurate non-invasive disease detection using cell-free DNA

**DOI:** 10.64898/2026.05.13.725033

**Authors:** Jingqi Li, Zhaoying Liu, Heng Zhang, Yuyang Zhang, Wei Li, Yumei Li

## Abstract

Plasma cell-free DNA (cfDNA) emerged as a promising non-invasive biomarker for cancers. However, reliable detection remains challenging due to the low abundance of tumor-derived cfDNA fragments and the dilution of informative methylation signals when aggregated into region-level features. Here, we propose a novel approach cfMIND, an efficient and robust machine-learning framework that leverages stratified read-level methylation signals to preserve rare cell-type–specific information and enhance detection sensitivity. By avoiding information loss inherent to conventional aggregation strategies, cfMIND enables more accurate and stable inference across diverse conditions. cfMIND is compatible with various cfDNA methylation sequencing technologies and cancer types. Across multiple cancer datasets (n = 868), cfMIND achieves high performance (AUROC up to 0.966) and maintains strong accuracy even at ultra-low sequencing depth (0.2×) and in early-stage cancers. Notably, cfMIND demonstrates exceptional robustness, generalizing effectively across cohorts and platforms without the need for model retraining. These results highlight its potential utility in heterogeneous experimental and clinically relevant settings. Beyond cancer detection, cfMIND is readily extendable to non-malignant diseases, as demonstrated by its ability to capture disease-associated methylation alterations in amyotrophic lateral sclerosis (ALS). Functional investigations on cfMIND-identified features further reveal enrichment in key regulatory regions implicated in disease pathogenesis and recapitulate tissue- and single-cell-level methylation and transcriptional programs underlying tumor biology. Collectively, cfMIND represents a significant advancement in the field, offering a broadly applicable, functionally interpretable, and high-resolution framework for non-invasive disease detection.

## Introduction

Liquid biopsy is a non-invasive diagnostic approach that analyzes biomarkers in body fluids (e.g., blood) to detect diseases, particularly cancer^1^. Among these biomarkers, cell-free DNA (cfDNA), which is predominantly double-stranded DNA fragments released by dying cells from various tissues, carries valuable genetic^2^, fragmentomic^3,4^, and epigenetic information^5–7^. In healthy individuals, the majority of cfDNA originates from blood cells^8^. However, during the progression and treatment of various diseases, particularly cancer, higher amounts of cfDNA derived from affected tissues (e.g., tumor tissue) can be detected in the plasma^9^, making it a promising biomarker for early detection and real-time monitoring of disease. Among various cfDNA-derived features, DNA methylation has emerged as a promising biomarker for early cancer detection, as aberrant DNA methylation has been frequently reported in cancer cells and can occur early in tumorigenesis^10,11^. Notably, cfDNA methylation has been reported to outperform other features, such as copy number alterations and fragment length profile^12^. Moreover, methylation alterations are detectable in cfDNA even when the patients have no significant clinical symptoms^13,14^, underscoring its potential for highly sensitive, non-invasive early cancer detection.

Nevertheless, since tumor-derived cfDNA represents only a small fraction of total cfDNA^15^, especially in early-stage cancer patients, accurately and sensitively detecting cancer-specific methylation signals against the overwhelming background of non-tumor cfDNA remains a significant technical challenge. To date, several studies have attempted to address this issue by leveraging DNA methylation profiles^16–18^. They depend heavily on matched tumor tissue and normal plasma samples to identify cancer-specific markers and train models, which is impractical in real-world clinical settings and susceptible to dataset-specific bias. Moreover, extending these methods to diverse cancer types requires labor-intensive manual marker selection or re-tuning, limiting scalability and reproducibility. In addition, the deep-learning based methods require substantial computational resources for model training, posing barriers to broader adoption. These constraints underscore the urgent need for a robust, efficient, and clinically applicable method that enables accurate cancer detection from cfDNA methylation profiles without relying on prior tissue data or intensive computational overhead.

To overcome these limitations, we developed cfMIND (*<u>cf</u>*DNA *<u>M</u>*ethylation signals of *<u>IN</u>*dividual read for disease *<u>D</u>*etection), a machine learning framework that leverages stratified read-level methylation signals from plasma cfDNA for sensitive and accurate disease detection. Unlike existing approaches constrained by tissue requirements or cancer-type specificity (**Supplementary Table 1**), cfMIND is compatible with diverse cfDNA methylation sequencing technologies, does not require matched tissue data, and readily generalizes across multiple cancer types. Its flexible design also enables direct application to non-cancerous conditions, broadening its clinical utility. Beyond classification performance, cfMIND uncovers biologically meaningful features that highlight key regulatory regions implicated in cancer pathogenesis and is concordant with tissue- and single-cell-level methylation transcriptional programs, offering both diagnostic power and mechanistic insight. Together, these advantages position cfMIND as a generalizable, clinically applicable, and biologically informative solution for cfDNA-based disease detection.

## Results

### cfMIND overview

The primary goal of cfMIND is to improve cancer diagnosis by exploiting read-level methylation information in plasma cfDNA, explicitly accounting for the heterogeneity of cancer cells and ensuring computational efficiency. Traditional DNA methylation analysis usually relies on the average methylation ratio of CpG sites across a genomic region. However, such averaging often obscures subtle signals from the small proportion of cfDNA derived from cancer cells: for example, if only 5% of cfDNA fragments originate from heterogeneous cancer cells (some hypermethylated, others hypomethylated), the overall average methylation may appear indistinguishable between cancer patients and healthy controls (**Fig. 1A).** To capture this signal in an efficient way, cfMIND introduces a read-stratification strategy. Specifically, the average methylation value of all CpG sites in a read (*r̅*) is discretized into five methylation levels by being rounded to the closest value in {0,0.25,0.5,0.75,1}. Then the read count for each level on a genomic region is summed and normalized to obtain five features {*M*_0′, *M*_0.25′, *M*_0.5′, *M*_0.75′, *M*_1′} (**Methods**). This encoding preserves the distributional heterogeneity of methylation patterns within a region (**Fig. 1A**), enabling detection of cancer-specific deviations. Using 500-bp genomic regions which can capture over 90% of CpGs in the human genome after retaining regions with more than three CpGs (**Supplementary Fig. 1**), cfMIND first extracts five-level features for each region and then applies statistical testing to identify disease-associated features, followed by Boruta-based feature selection to retain high-confidence signals (**Fig. 1B**). These features are then integrated into an XGBoost classifier, which outputs individualized disease probability scores (**Fig. 1B**). This design uniquely leverages read-level stratification of cfDNA methylation to sensitively capture rare, heterogeneous cancer signals that are invisible to conventional averaging approaches.

**Fig. 1.**
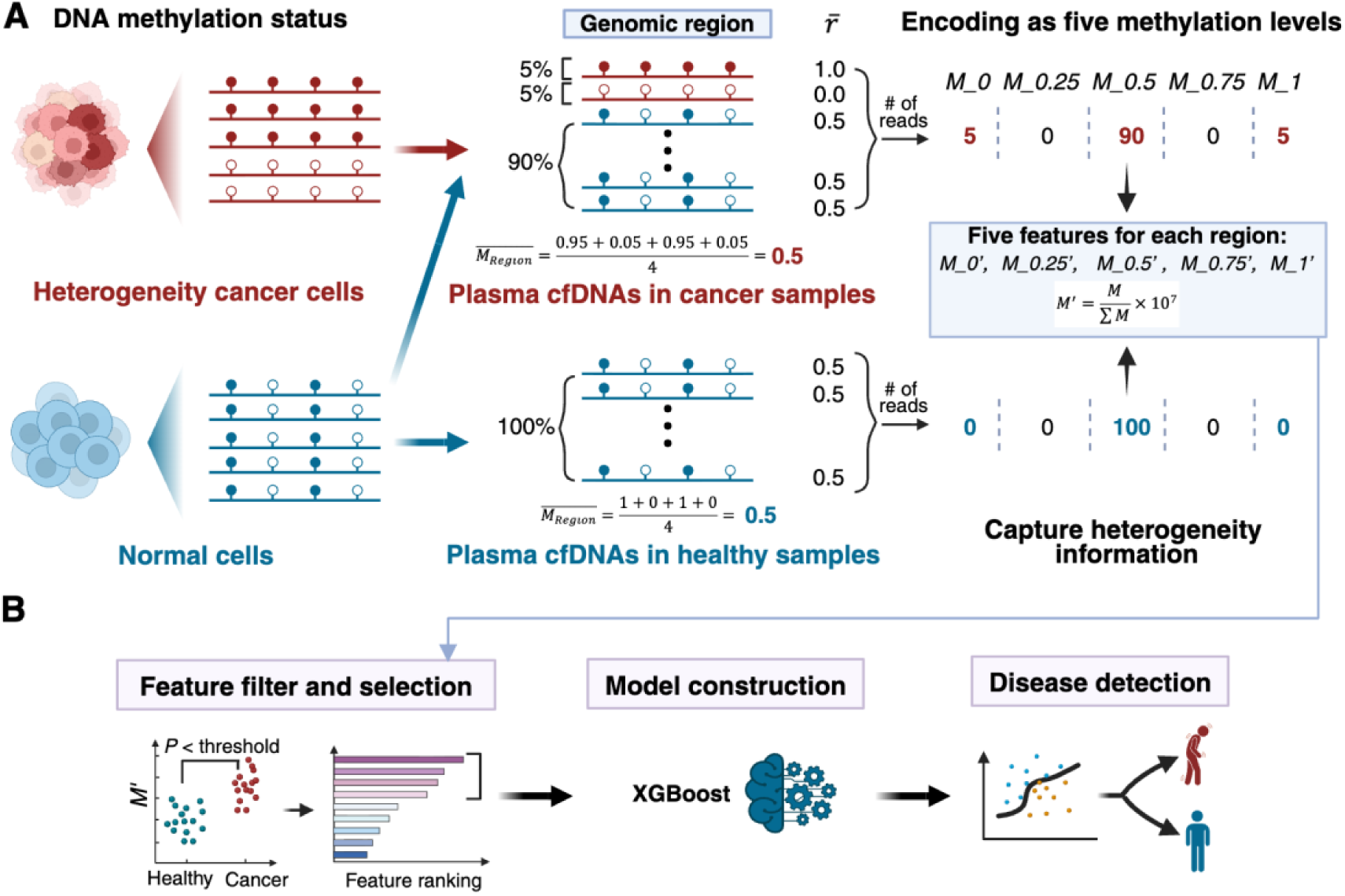
Overview of the cfMIND framework. **(A)** Rationale of the read-stratification strategy. In this example, cfDNAs from cancer cells show heterogeneous methylation patterns, with some fragments fully methylated (average methylation ratio *M̅* = 1) and others unmethylated (*M̅* = 0), while cfDNAs from normal cells are partially methylated ( *M̅* = 0.5). When averaged across the genomic region, both healthy samples (100% normal cfDNA) and cancer samples (10% cancer-derived cfDNA + 90% normal cfDNA) yield the same mean methylation ratio (0.5), making them indistinguishable. By discretizing the average methylation of each read *r̅* in the genomic region into five levels, cfMIND captures the distributional differences and reveals distinct features (e.g., *M*_0′, *M*_0.5′, *M*_1′) that reflect cancer-specific heterogeneity. **(B)** Overview of the cfMIND workflow. The five-level features extracted from each genomic region are first subjected to statistical filtering and Boruta-based feature selection. The resulting high-confidence features are then used to train an XGBoost classifier, which predicts the probability of disease for each sample.

### cfMIND demonstrates superior performance across various cfDNA methylation sequencing technologies and cancer types

As an initial evaluation, we assessed the performance of cfMIND in distinguishing cancer patients from non-cancer controls. Given the rapid advancement of high-throughput cfDNA methylation sequencing technologies—including the standard whole-genome bisulfite sequencing (WGBS)^19,20^, cost-effective cfMethyl-Seq^21^, newly-developed bisulfite-free methods enzymatic methyl sequencing (EM-seq)^22^, and TET-assisted pyridine borane sequencing (cfTAPS)^23^ (**Supplementary Table 2**)—we evaluated whether cfMIND could maintain robust performance across platforms. For liver hepatocellular carcinoma (LIHC) detection, we utilized four independent cohorts corresponding to these sequencing platforms: WGBS (n = 56; 24 LIHC and 32 healthy controls), cfMethyl-Seq (n = 223; 30 LIHC and 193 healthy controls), EM-seq (n = 200; 100 LIHC and 100 healthy controls), and cfTAPS (n = 51; 21 LIHC and 30 healthy controls). Using samples spanning both early- and late-stage disease, cfMIND achieved consistently high performance on all four technologies, with areas under the receiver operating characteristic curve (AUROC) ranging from 0.941 to 0.963 (**Fig. 2A**).

**Fig. 2.**
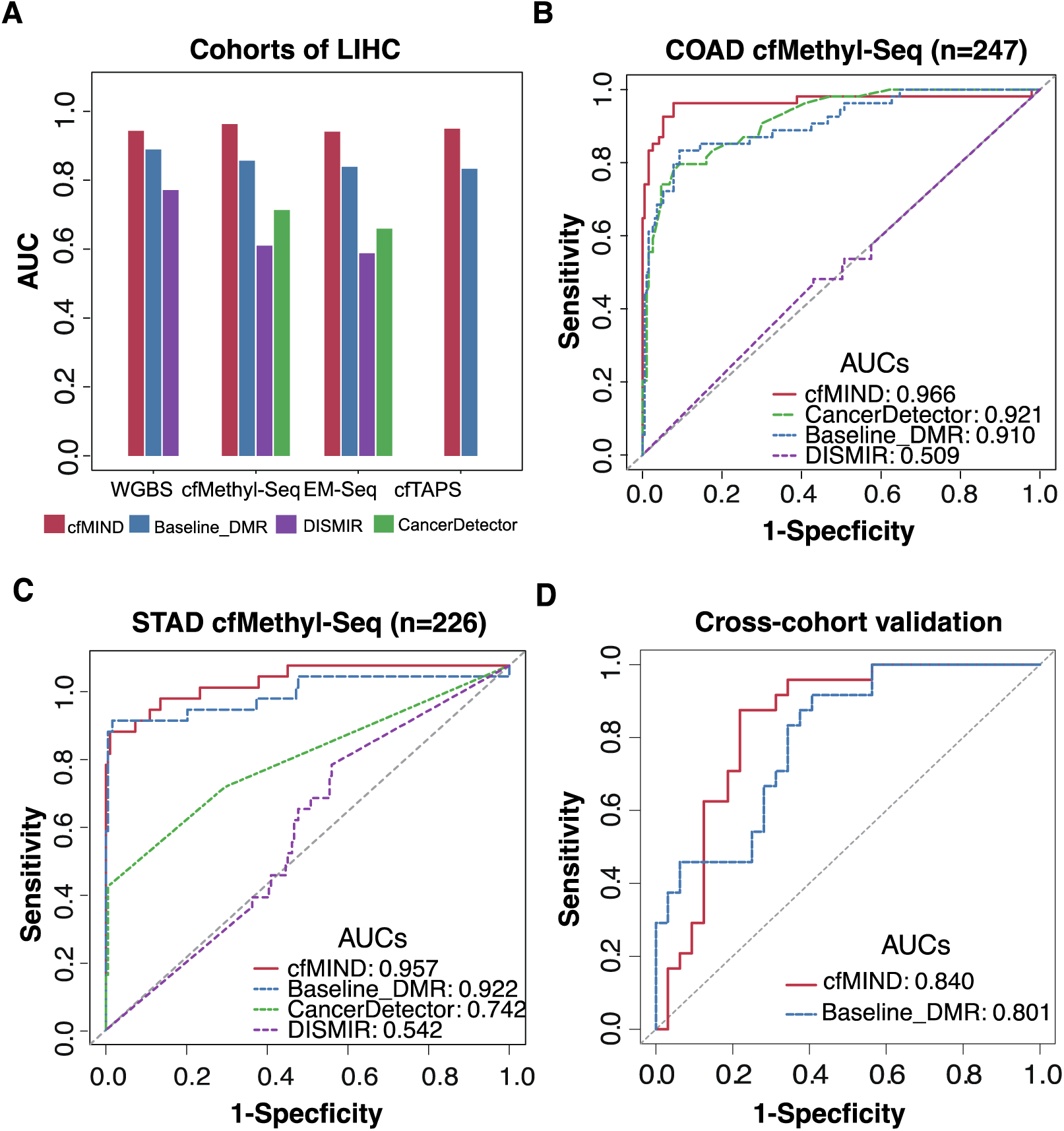
Performance of cfMIND in cancer detection across multiple sequencing technologies, cancer types, and cohorts. **(A)** Areas under the receiver operating characteristic curves for LIHC detection using cfMIND, Baseline_DMR, CancerDetector, and DISMIR for cohorts across four cfDNA methylation sequencing technologies: WGBS (n = 56; 24 LIHC and 32 healthy), cfMethyl-Seq (n = 223; 30 LIHC and 193 healthy), EM-seq (n = 200; 100 LIHC and 100 healthy), and cfTAPS (n = 51; 21 LIHC and 30 healthy). **(B-C)** ROC curves for COAD **(B)** and STAD **(C)** detection using cfMIND and comparator methods. **(D)** ROC curve showing the performance of cfMIND and Baseline DMR in an independent cross-cohort external validation WGBS cohort (n = 54; 17 LIHC and 37 healthy controls). All ROC curves were based on leave-one-out cross-validation analysis, and the number of samples for each dataset was indicated in parentheses.

To benchmark cfMIND, we compared its performance with existing cfDNA methylation-based cancer detection methods, including CancerDetector^16^, DISMIR^17^, and a baseline model using differentially methylated regions (Baseline_DMR; see **Methods**). Notably, existing tools have platform restrictions: CancerDetector requires paired-end sequencing data and is therefore not applicable to single-end datasets, the WGBS dataset. Additionally, CancerDetector and DISMIR are designed to process bisulfite-converted sequencing reads, rendering them incompatible with cfTAPS data due to its distinct methylation conversion technique. Despite these methodological differences, cfMIND consistently outperformed the comparator methods across platforms. It achieved the highest AUROC and sensitivity at 90% specificity in leave-one-out cross-validation (LOOCV) analyses **(Fig. 2A; Supplementary Table 3, and Supplementary Fig. 2A-D)**, demonstrating its robustness and adaptability to diverse sequencing technologies. Beyond LIHC, we evaluated the generalizability of cfMIND across cancer types. When applied to additional datasets, including a COAD cohort (n=247) and a STAD cohort (n=226) profiled via cfMethyl-Seq, cfMIND maintained top performance among the tested methods (**Fig. 2B-C; Supplementary Table 3, and Supplementary Fig. 2E-F)**, supporting its applicability across tumor types within a consistent experimental framework.

To further assess cross-cohort and cross-platform generalizability, we performed an external validation analysis in which a model trained exclusively on the cfMethyl-Seq cohort (n = 223), which yielded the highest internal performance, was applied without retraining to an independent WGBS cohort (n=56) generated using a distinct sequencing protocol. Under this setting, although cfMIND showed a measurable performance decrease relative to cross-validation, it still retained a relatively strong discriminative performance (AUROC = 0.84; **Fig. 2D**), despite differences in library preparation, sequencing chemistry, and data generation. Notably, only cfMIND and the Baseline_DMR were evaluated in this external validation setting. CancerDetector and DISMIR were not included because they are not compatible with cross-platform or cross-library applications: CancerDetector requires paired-end sequencing, and both methods are restricted to bisulfite-converted data, precluding direct application to single-end WGBS or cfTAPS datasets without substantial modification.

Overall, while performance decreases under external validation, cfMIND demonstrates stable and generalizable behavior across heterogeneous technical settings. Together with its consistent performance across datasets and platforms without retraining, these results support that cfMIND captures transferable methylation patterns rather than overfitting to dataset-specific features.

### cfMIND enables accurate early-stage cancer detection, cancer subtyping, and multi-classification

We next evaluated cfMIND in more challenging yet clinically critical scenarios, including early-stage detection and cancer subtyping. Detecting early-stage cancers is difficult because tumors shed only a small fraction of cfDNA into the bloodstream, often less than 1–5% of the total cfDNA^24^, making tumor-derived methylation signals rare and easily masked by the overwhelming background of cfDNA from normal tissues. In addition, early-stage tumors are often more heterogeneous^25,26^, further diluting the detectable methylation alterations. By design, cfMIND can capture these subtle and heterogeneous patterns using the read-stratification strategy. Across four cancer types (COAD, LUAD, LUSC, and STAD) from the cfMethyl-Seq dataset, cfMIND achieved great performance in detecting late-stage cancers (mean AUROC: 0.955 to 0.990), confirming its ability to capture strong tumor-derived cfDNA signals when disease burden is high (**Fig. 3A; Supplementary Table 4**). More importantly, cfMIND maintained robust performance in early-stage cancers, achieving AUROC values of 0.881 to 0.907 (**Fig. 3A; Supplementary Table 4**), a setting where cfDNA is scarce and signals are easily masked by background noise, underscoring its potential value for early cancer detection and population-level screening.

**Fig. 3.**
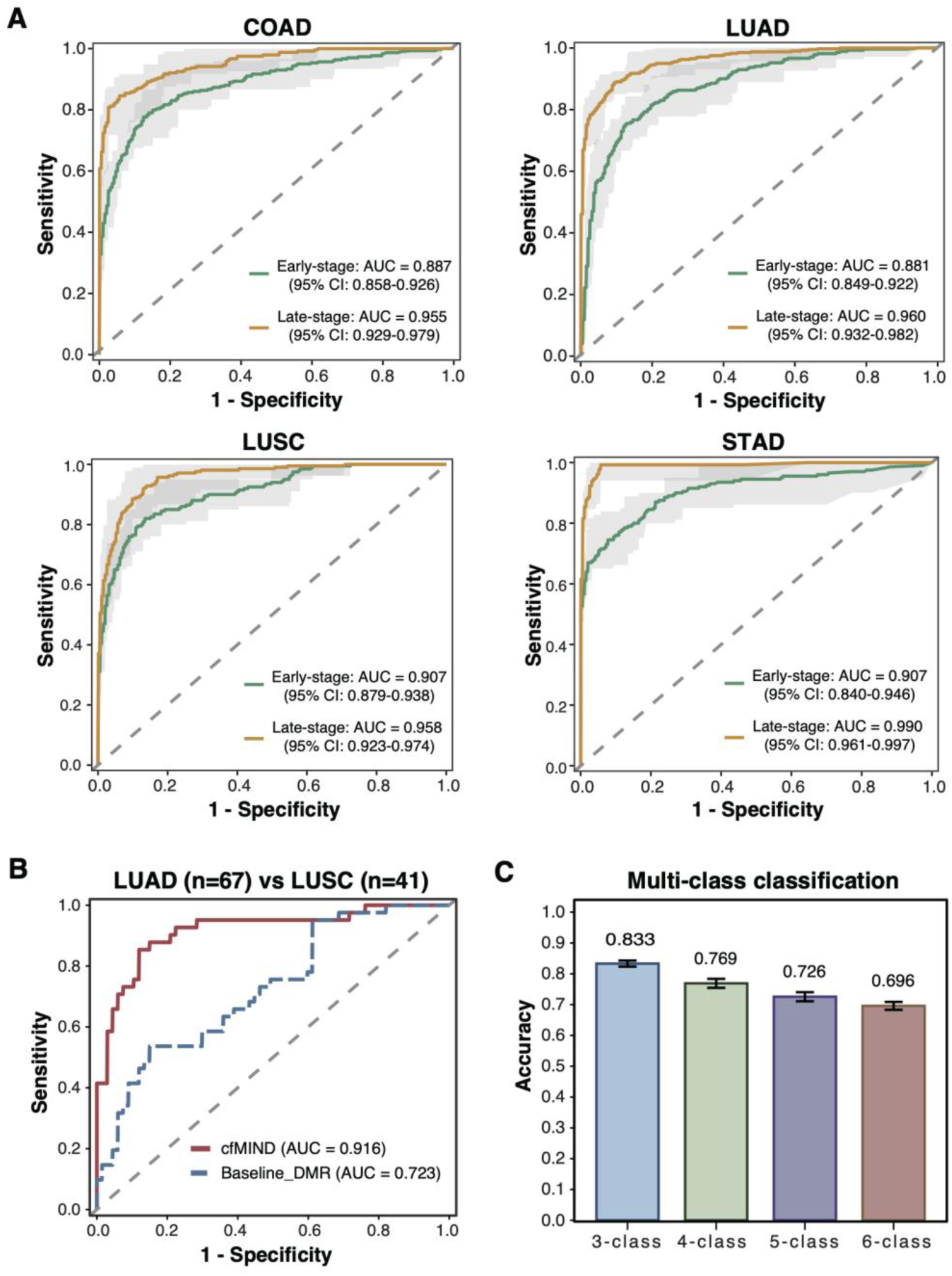
Performance of cfMIND in early cancer detection, subtype classification, and multi-class tasks. **(A)** ROC curves for early- and late- stage cancer detection using samples from the cfMethyl-Seq dataset. Solid lines represent mean performance, and shaded areas indicate 95% confidence intervals from 10 repetitions of 5-fold cross-validation. **(B)** ROC curves for distinguishing LUAD and LUSC using cfMIND and Baseline_DMR from cfMethyl-Seq dataset. AUCs are calculated with leave-one-out cross-validation. **(C)** Bar plots showing the average classification accuracy of cfMIND in multi-class tasks (3-6 classes) constructed by sequentially adding cancer types to healthy controls from the cfMethyl-Seq dataset. Error bars represent standard deviations across 10 repetitions of 5-fold cross-validation.

We next assessed cfMIND’s ability to classify cancer subtypes. For lung cancer, subtype distinction is clinically significant, as lung adenocarcinoma (LUAD) and lung squamous cell carcinoma (LUSC) differ in molecular features, treatment responses, and patient outcomes, yet often present with overlapping clinical profiles^27–29^. Leveraging cfDNA methylation signatures from the cfMethyl-Seq dataset (comprising 67 LUAD and 41 LUSC samples), cfMIND distinguished LUAD from LUSC with an AUROC of 0.916, outperforming the Baseline_DMR method (**Fig. 3B; Supplementary Table 5**). Such non-invasive stratification not only supports precision diagnosis but also guides treatment selection and monitoring, ultimately improving patient outcomes.

Finally, we extended cfMIND to multi-class classification tasks involving three to six classes using the cfMethyl-Seq dataset. Each task included the healthy control group together with multiple cancer types, with classes added sequentially in descending order of sample size to ensure stable training. Specifically, the 3-class task comprised healthy controls, LUAD, and COAD; the 4-class task added LUSC; the 5-class task further included STAD; and the 6-class task encompassed all groups (healthy controls, LUAD, COAD, LUSC, STAD, and LIHC). Under this design, cfMIND achieved >80% accuracy in the 3-class setting and maintained ∼70% accuracy in the 6-class setting (**Fig. 3C**). We observed a gradual decrease in accuracy as the number of classes increased. This trend is expected given the increasing classification complexity and greater overlap in methylation profiles among cancer types. The relatively smooth performance decline, rather than abrupt drops associated with specific class additions, suggests that classification difficulty is driven primarily by the expanding label space rather than any single cancer type.

### Robustness and efficiency of cfMIND across clinically realistic scenarios

As cfMIND is ultimately designed for clinical translation, we next evaluated its robustness under conditions that better mimic real-world applications. To account for cost constraints in clinical practice, we first tested cfMIND on a targeted bisulfite sequencing dataset (targeted regions ≈ 30 Mb; 1% of the human genome). cfMIND again delivered the best performance for LIHC detection on this platform (**Fig. 4A; Supplementary Table 3; Supplementary Fig. 3A**). We then examined the effect of sequencing depth using LUAD samples from the cfMethyl-Seq dataset which has the largest sample size (n = 260). The original dataset was down-sampled to simulate different sequencing depth, ranging from 0.2× to 20×, with each condition repeated five times. cfMIND achieved an average AUROC of 0.736 even at an ultra-low sequencing depth of 0.2×, and performance improved rapidly, reaching 0.897 at 1× and stabilizing at 0.958 from 5× upwards (**Fig. 4B**). Notably, as depth decreased, the number of features declined but the retained features remained high performance, reflecting cfMIND’s efficiency in capturing disease signals despite sparse data (**Supplementary Fig. 3B**). Together, these results highlight cfMIND’s robustness to sequencing noise and data sparsity, supporting its suitability for clinical settings where cost or sample constraints may limit sequencing depth.

**Fig. 4.**
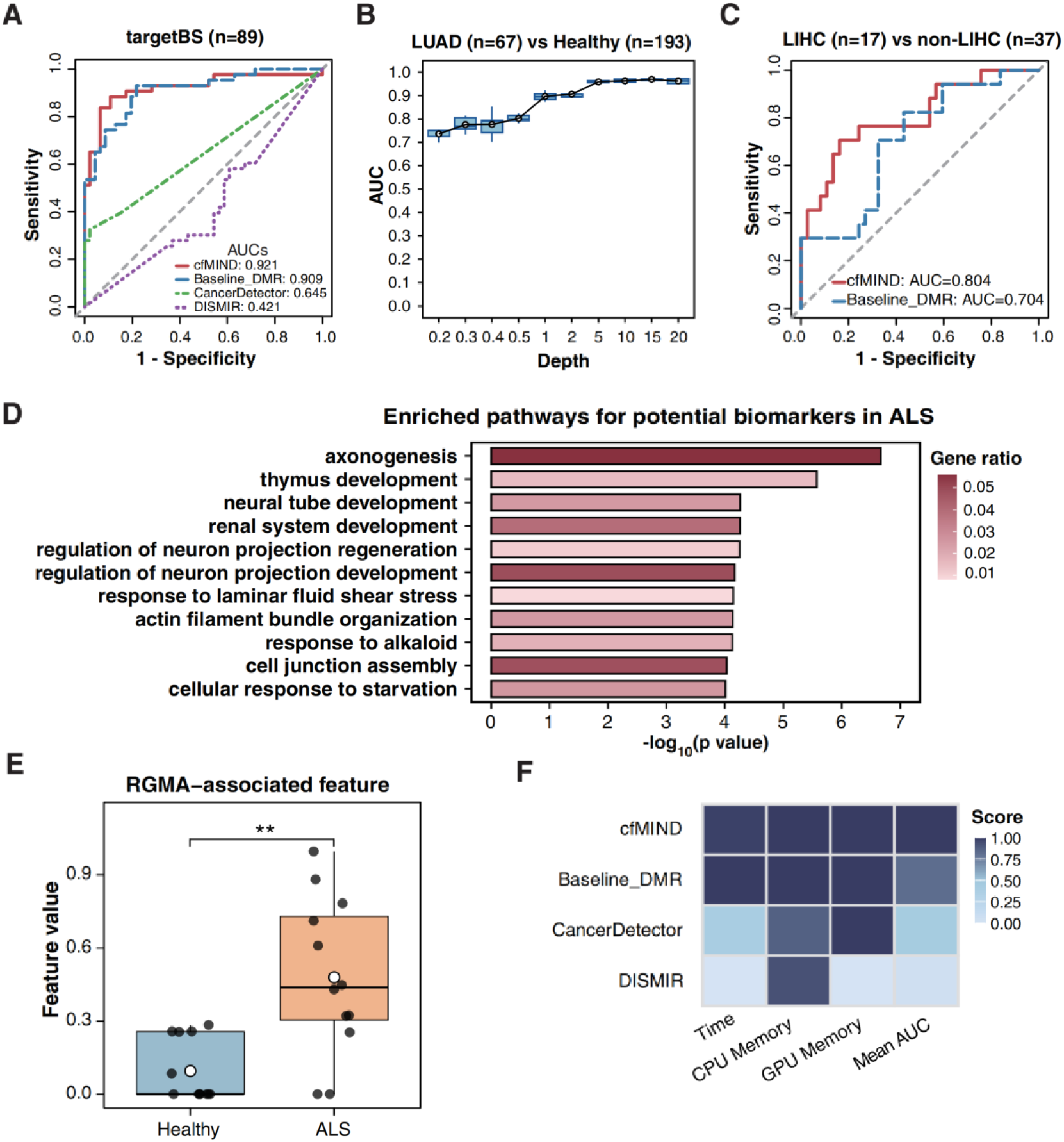
Performance and utility of cfMIND across diverse clinical scenarios. **(A)** ROC curves for detecting LIHC in a targeted bisulfite sequencing dataset (targetBS). **(B)** Boxplots showing AUC values of LIHC detection at varying sequencing depths, assessed by LOOCV with five replicates per depth via subsampling. White dots represent the mean AUC at each depth. **(C)** Comparisons of ROC curves for distinguishing LIHC (n=17) from non-LIHC including cirrhosis (n=17), hepatitis (n=17), and healthy controls (n=3) between cfMIND and Baseline DMR evaluated by LOOCV. **(D)** Barplots showing significantly enriched pathways (adjusted p-value < 0.05) from functional enrichment analysis of genes linked to the top 2,000 cfMIND features ranked by feature importance for an ALS cohort (n = 24; 12 ALS and 12 healthy controls) profiled by WGBS. **(E)** Boxplots showing the feature values for RGMA-associated feature in ALS patients and healthy controls for the ALS cohort. Wilcoxon rank-sum test was used to calculate p-value between the two groups. **(F)** Heatmap summarizing the performance of cfMIND and comparator methods across three usability metrics and the mean AUC values for all datasets evaluated in Fig. 2. Each metric was normalized to a 0–1 scale, with higher values indicating better performance. See the “Methods” section for details on metric definitions and normalization procedures.

Furthermore, we tested cfMIND in more challenging classification scenarios using an additional WGBS cohort (n=54, consisting of 17 LIHC, 17 cirrhosis, 17 hepatitis, and 3 healthy controls). Distinguishing LIHC from benign liver diseases, such as hepatitis and cirrhosis, remains difficult; nevertheless, cfMIND achieved an AUROC value of 0.804. (**Fig. 4C; Supplementary Table 6**). In contrast, existing methods, such as CancerDetector and DISMIR are primarily designed to distinguish cancer from normal samples, and thus not applicable to this task. Moreover, because these approaches depend on tumor tissue samples for marker discovery, they obtained inconsistent marker set and unstable performance across datasets (**Supplementary Fig. 4**), reflecting their intrinsic limitations and susceptibility to dataset-specific bias.

Beyond cancer, non-invasive detection and monitoring is also essential for other fatal diseases. One example is ALS, a progressive and fatal neurodegenerative disorder affecting motor neurons. The potential of cfDNA for early ALS detection has attracted growing interest in neuroscience^30^. Using a cfDNA WGBS dataset of 12 ALS patients and 12 age-matched controls, we evaluated whether cfMIND could capture disease-relevant signals. From the top 2,000 features ranked by cfMIND’s feature importance, the associated genes were significantly enriched in neuronal processes, particularly axonogenesis (**Fig. 4D**), a biological pathway central to ALS pathogenesis^31^. Among the genes involved in axonogenesis, *RGMA*—previously reported to be abnormally elevated in ALS and to promote axonal damage^32^—showed significant feature differences between ALS patients and controls (**Fig. 4E; Supplementary Fig. 5**). These findings highlight cfMIND’s ability to uncover pathogenesis-associated biomarkers in fatal neurodegenerative diseases, extending its utility beyond cancer.

Finally, we evaluated the computational efficiency of cfMIND. Unsurprisingly, cfMIND and Baseline_DMR ran substantially faster than the other three methods, including a probabilistic framework and two deep learning models, in leave-one-out cross-validation analyses (**Supplementary Fig. 6A)**. Notably, cfMIND required minimal memory—only 272 MiB—due to its streamlined modeling strategy, enabling execution on standard personal computers (**Supplementary Fig. 6B),** while deep learning methods demanded far greater computational resources. When integrating accuracy and efficiency into a composite performance score (see **Methods**), cfMIND ranked the highest, combining superior AUROC with minimal computational cost (**Fig. 4F**).

### cfMIND identifies biologically meaningful and robustly differential features

To investigate the characteristics and biological relevance of the features prioritized by cfMIND, we focused on COAD, one of the most prevalent and lethal cancers worldwide^33^. Using the COAD cfMethyl-Seq dataset, we first examined the five-level features included in the final classifier. These features were distributed across diverse genomic regions (**Fig. 5A**), indicating that cfMIND captures a broad spectrum of informative loci. When compared with the features used in the Baseline_DMR model, only 11.56% of genomic regions overlapped (**Fig. 5B**), demonstrating that cfMIND identifies additional disease-relevant signals beyond those detected by conventional DMR approaches. For the overlapping regions, cfMIND’s five-level representation further amplified the differences between cancer and control samples (**Fig. 5B**), highlighting its enhanced sensitivity to subtle methylation alterations.

**Fig. 5.**
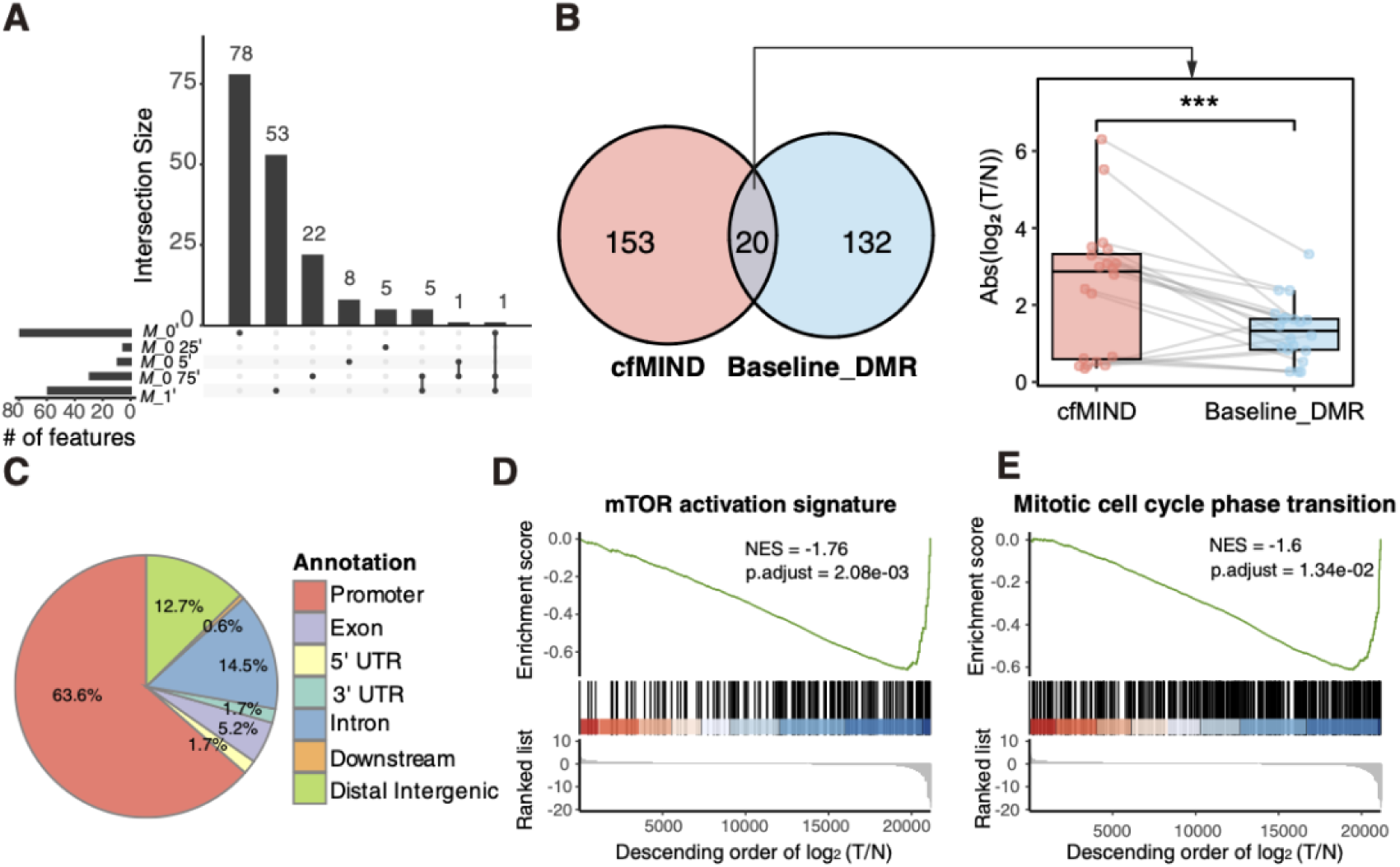
The characteristics of features identified by cfMIND. **(A)** Upset plots illustrate the genomic location intersections of the five-level features. **(B)** Overlap of features between cfMIND and Baseline_DMR models (**left panel**) and absolute feature fold change between tumor and normal cfDNA samples for the overlapped features (**right panel**). **(C)** Pie chart showing the genomic annotation of features identified by cfMIND. **(D-E)** Enrichment plot showing the GSEA results for mTOR activation signature **(D)** and mitotic cell cycle phase transition pathway **(E)**. The green curve represents the running enrichment score (ES) across the ranked gene list, with the peak valley indicating the minimum ES. The vertical black lines denote the positions of pathway genes within the ranked list, and the color gradient (red to blue) reflects gene-level correlation with the cfMIND feature values. The histogram displays log2-transferred fold change of feature values between tumor and normal samples. The normalized enrichment score (NES) and adjusted p-value (p.adjust) are shown in the plot.

Genomic annotation analysis revealed that most cfMIND-derived features were located in functional regions such as promoters, exons, and introns (**Fig. 5C**). To further assess their biological significance, we performed Gene Set Enrichment Analysis (GSEA) on genes associated with these features, identifying multiple gene sets significantly enriched for pathways known to drive cancer development (**Fig. 5D–E; Supplementary Fig. 7**). Notably, the mTOR signaling pathway—an established therapeutic target in colorectal cancer ^34–36^—was among the top enriched pathways (**Fig. 5D; Supplementary Table 7**). Another enriched gene set, mitotic cell-cycle phase transition, plays a central role in tumor progression by enabling uncontrolled proliferation^37,38^ (**Fig. 5E; Supplementary Table 8**). Together, these results demonstrate that cfMIND not only detects robust differential methylation signals but also highlights biologically meaningful features linked to cancer pathogenesis and potential therapeutic opportunities.

### cfMIND recapitulates the tumor tissue- and single-cell- level methylation and transcriptional alterations

As cfMIND was designed to capture the subtle signals from the small fraction of tumor-derived cfDNA, we next evaluated its ability to identify tumor-associated methylation alterations. Notably, in the COAD cfMethyl-Seq dataset, the majority of top-ranked cfMIND features were consistently recovered across cross-validation folds, indicating that the identified biological signals are robust rather than driven by model instability (**Supplementary Fig. 8**). We then categorized the top-ranked features into four groups based on differences in *M*_0′ and *M*_1′ between cancer patients and healthy individuals: *M_0.TvsN.down* (lower *M*_0′in COAD patients), *M_0.TvsN.up* (higher *M*_0′ in COAD patients), *M_1.TvsN.down* (lower *M*_1′ in COAD patients), and *M_1.TvsN.up* (higher *M*_1′in COAD patients) (**Methods; Fig. 6A; Supplementary Table 9-11**). Using tumor and normal tissue methylation array data (n=349; 311 COAD and 38 normal tissues) from The Cancer Genome Atlas (TCGA)^39^, and restricting the analysis to regions containing ≥ 3 CpGs and categories with ≥ 50 regions (**Supplementary Table 9**), we observed that regions in the *M_0.TvsN.down* and *M_1.TvsN.up* groups exhibited higher methylation level in tumor tissues, indicating that cfMIND recapitulates both downregulated low-methylation cfDNA fragments and upregulated high-methylation cfDNA fragments derived from tumor tissues (**Fig. 6B**). Given the well-established relationship between DNA methylation and transcriptional regulation^40,41^, we next assessed whether these cfMIND-derived methylation features corresponded to transcriptional alterations. Using TCGA COAD bulk RNA-seq data (n=524; 483 COAD and 41 normal tissues), genes whose promoters associated with hypermethylated regions showed reduced expression in tumors compared with normal tissues (**Fig. 6C**), consistent with the expected inverse relationship between promoter methylation and gene expression. This observation suggests that cfMIND-derived features effectively reflect tumor-associated transcriptional dysregulation.

**Fig. 6.**
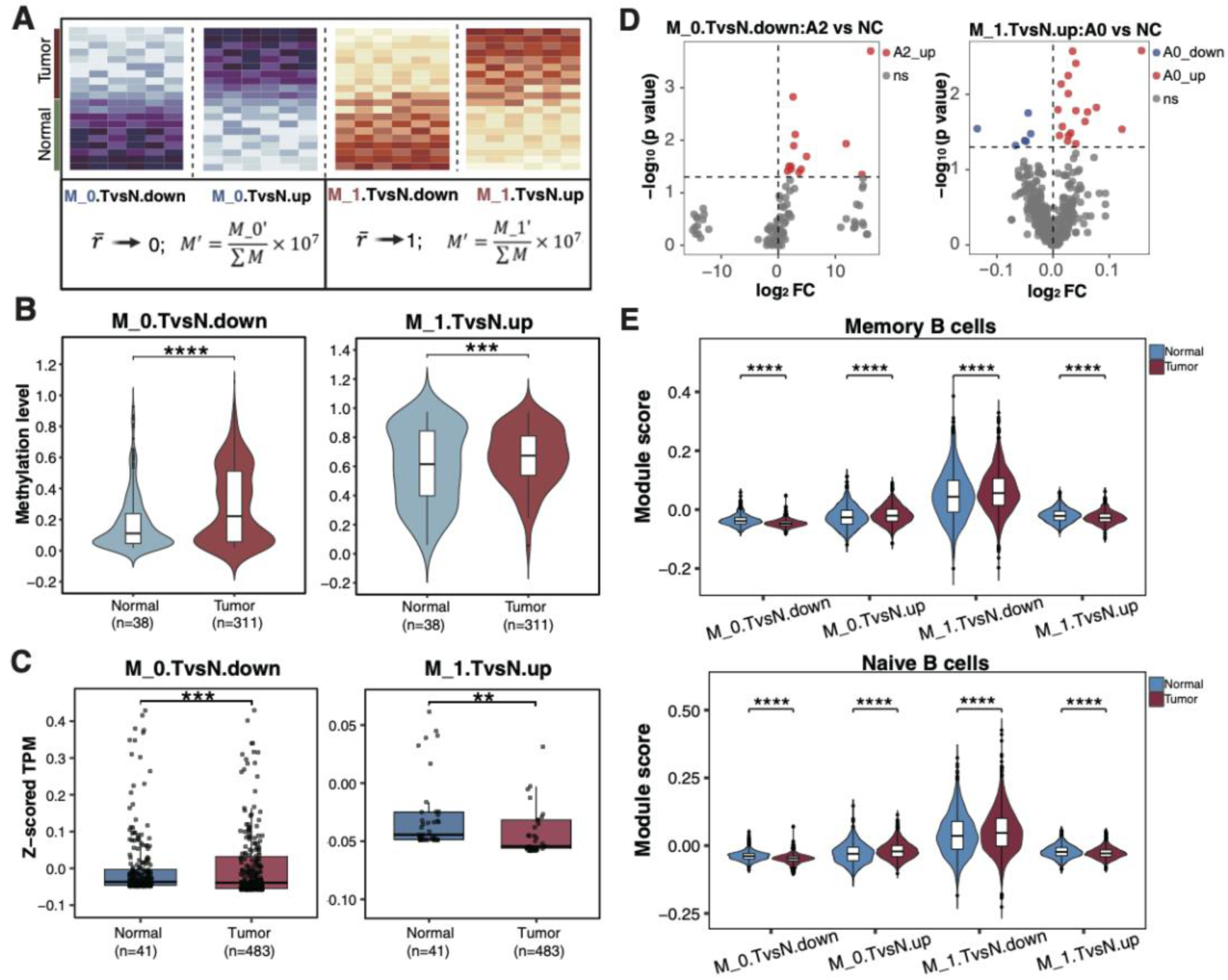
Characterization of cfMIND-derived features for tumor tissue and single-cell levels. **(A)** Diagram illustrating the feature regions with significant *M*_0′ (left) and *M*_1′ (right) differences between COAD and normal cfDNA samples. Features were categorized as four groups, and the color gradients represent the normalized feature value *M*′ computed from either *M*_0′ or *M*_1′. **(B)** Violin plots showing mean DNA methylation levels for *M_0.TvsN.down* (left panel) and *M_1.TvsN.up* (right panel) in normal and COAD tumor tissues using TCGA COAD methylation array data (n = 349; 311 COAD and 38 normal). The number of samples for each dataset was indicated in parentheses. **(C)** Boxplots showing expression level of genes whose promoter regions mapped to *M_0.TvsN.down* (left panel) and *M_1.TvsN.up* (right panel) using TCGA COAD bulk RNA-seq data (n = 524; 483 COAD and 41 normal). The number of samples for each dataset was indicated in parentheses. **(D)** Volcano plots showing differentially methylated features in *M_0.TvsN.down* (left panel) and *M_1.TvsN.up* (right panel) at the single-cell level using a single bisulfite sequencing dataset (CRC01) comprising 359 cells (343 COAD and 16 normal). Red dots indicate regions that are significantly hypermethylated in the COAD lineage, blue dots indicate regions that are hypomethylated, and gray dots represent regions without significant methylation changes. **(E)** Violin plots comparing meta-gene expression levels (module score) between normal and tumor memory B cells (top) and naive B cells (bottom) using integrated data from 6 independent single-cell RNA-seq datasets (total 57 samples; 34 COAD and 23 normal). Genes whose promoter regions mapped to features in the four groups were included in the analysis. P-values were determined using one-sided Wilcoxon rank-sum test. **P < 0.05, ***P < 0.001, ****P < 0.001.

To further investigate the cellular origin of these cfMIND features, we conducted single-cell level analyses using both single-cell methylation^42^ and single-cell RNA sequencing data^43–47^. Using single-cell methylation data from a colorectal cancer patient (comprising 343 COAD cells and 16 normal cells), we compared methylation changes of *M_0.TvsN.down* and *M_1.TvsN.up* regions across 12 sub-lineages identified from tumor and the adjacent normal colon tissue. Several sub-lineages exhibited significantly higher levels of upregulated features compared with adjacent normal colon cells (**Fig. 6D; Supplementary Fig. 9A; Supplementary Table 10**), indicating cfMIND recapitulates cell-type–specific methylation signatures contributing to tumor–normal discrimination in cfDNA. Consistently, single-cell RNA-seq analysis (integrating 6

independent datasets with a total of 34 COAD and 23 normal samples) showed significantly reduced expression of genes associated with hypermethylated regions (*M_0.TvsN.down* and *M_1.TvsN.up*) in immune cells such as Naïve and memory B cells, while genes linked to hypomethylated regions (*M_0.TvsN.up* and *M_1.TvsN.down*) were upregulated in tumors (**Fig. 6E; Supplementary Fig. 9B; Supplementary Table 11**). These results suggest that immune cell–associated methylation and transcriptional programs may contribute meaningfully to cfDNA-based tumor–normal discrimination, consistent with the known role of tumor–immune interactions in colorectal cancer^48–50^.

## Discussion

In this study, we present cfMIND, a machine-learning framework that enables sensitive detection of tumor-derived signals from plasma cfDNA by leveraging stratified read-level DNA methylation patterns. By modeling methylation states within individual cfDNA fragments rather than relying solely on regional or bulk-level averages, cfMIND addresses a central challenge in liquid biopsy analysis: the accurate identification of rare tumor-derived DNA molecules against a dominant background of cfDNA released from normal tissues. Across multiple datasets, cancer types, and sequencing platforms, cfMIND consistently achieved highest diagnostic performance, even at substantially reduced sequencing depths, underscoring its robustness and translational potential.

A key conceptual advance of cfMIND lies in its use of read-level methylation heterogeneity to capture subtle tumor-associated signals. Traditional cfDNA methylation analyses typically aggregate methylation information across genomic regions^51,52^, which can obscure informative fragment-level patterns, particularly when tumor DNA fractions are low. By stratifying cfDNA reads based on their internal methylation configurations, cfMIND effectively amplifies tumor-specific epigenetic signatures without requiring explicit tumor fraction estimation or matched tissue references. This design enables high sensitivity while maintaining compatibility with diverse cfDNA methylation sequencing technologies, including both targeted and genome-wide assays.

Beyond performance metrics, cfMIND provides biologically meaningful insights into the origins and functional relevance of its predictive features. Integrative analyses using TCGA tumor methylation and transcriptome data demonstrated that cfMIND-identified regions reflect bona fide tumor-associated epigenetic alterations and are concordant with transcriptional dysregulation in cancer tissues. Furthermore, single-cell–level analyses revealed that these features originate from specific cellular subpopulations within the tumor microenvironment, including immune cell lineages. These findings suggest that cfMIND captures a composite signal arising not only from malignant epithelial cells but also from tumor-associated immune components, consistent with emerging evidence that cfDNA reflects the broader tissue ecosystem rather than tumor cells alone^53^. This property may be particularly advantageous for early detection, where immune remodeling often precedes overt tumor expansion^54,55^.

Another notable strength of cfMIND is its extensibility beyond oncology. Its successful application to detecting methylation alterations associated with ALS illustrates that the framework is not limited to cancer-specific signals but can be generalized to other diseases characterized by subtle cfDNA methylation changes. This expands the potential utility of cfMIND to a wide range of pathological conditions, including neurodegenerative^30,56^, inflammatory^57^, and autoimmune diseases^58^, where non-invasive biomarkers are critically needed.

Despite these strengths, several limitations warrant consideration. First, while cfMIND captures tumor- and immune-associated signals, the precise quantitative contribution of different tissue and cell types to the cfMIND model remains to be fully characterized. Future studies integrating high-resolution tissue-of-origin deconvolution may further enhance interpretability. Second, although cfMIND performed robustly across retrospective datasets, prospective validation in large, clinically representative cohorts—particularly in early-stage and pre-diagnostic settings—will be essential to establish its clinical utility. Finally, while the current study focused on DNA methylation, integrating additional cfDNA features such as fragmentomics^59^ or nucleosome positioning^60^ may further improve sensitivity and disease specificity.

## Methods

### cfMIND overview

cfMIND is a read-level cfDNA methylation-based framework designed to improve the sensitivity and robustness of disease detection by capturing methylation heterogeneity. The cfMIND workflow comprises two modules: **(i)** extraction of read-level features across five discrete methylation levels; and **(ii)** construction of a disease detection machine learning model. For feature extraction, cfMIND first calculated the average methylation ratio (*r̅*) for each read with ≥ 3 CpGs. *r̅* was defined as the number of methylated CpGs divided by the total number of CpGs covered by the read and rounded to the nearest value in {0,0.25,0.5,0.75,1}. Next, the genome was divided into non-overlapping 500 bp bins, retaining only regions containing at least three CpGs and with an average sequencing coverage above a predefined threshold (default: 20). For each region, cfMIND counted the number of reads assigned to each methylation levels, yielding five features (*M*_0, *M*_0.25, *M*_0.5, *M*_0.75, *M*_1). To account for differences in sequencing depth across samples, features were normalized by the total read count within each sample according to 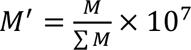, resulting in the final normalized features (*M*_0′, *M*_0.25′, *M*_0.5′, *M*_0.75′, *M*_1′) for downstream analysis. For model construction, cfMIND first performed two-sided t-tests on alsl features to identify those with significant differences between groups (p < 0.01). Informative features were then selected using the Boruta algorithm implemented in the Boruta R package (v8.0.0; *pValue=0.01, mcAdj =TRUE, maxRuns=500*). The number of final features selected for each dataset is summarized in **Supplementary Table 12**. Finally, disease classification models were trained using eXtreme Gradient Boosting (XGBoost; xgboost R package v1.7.7.1), supporting both binary and multi-class prediction tasks. Cross-validation was applied throughout model training to prevent overfitting.

### Processing of methylation sequencing data for cfDNA and solid tissues

In total, seven cfDNA methylation sequencing datasets and one tissue methylation sequencing dataset were included in this study. To ensure a comprehensive evaluation across diverse sequencing platforms and clinical scenarios, the cfDNA cohorts comprised the following: (1) WGBS cohort (32 healthy controls, 24 LIHC); (2) cfMethyl-Seq cohort representing multiple cancer types (193 healthy controls, 30 LIHC, 54 COAD, 67 LUAD, 41 LUSC, 33 STAD); (3) EM-seq cohort (100 healthy controls, 100 LIHC); (4) cfTAPS cohort (30 healthy controls, 21 LIHC); (5) targeted bisulfite sequencing (targetBS) cohort (46 healthy controls, 43 LIHC); (6) an additional WGBS cohort of 54 samples for benign disease confounding analysis (3 healthy controls, 17 LIHC, 17 cirrhosis, 17 hepatitis); and (7) 24 WGBS samples for non-cancerous disease evaluation (12 healthy controls, 12 ALS). Furthermore, a solid tissue Reduced Representation Bisulfite Sequencing (RRBS) dataset containing 68 tumor/normal pairs (comprising 53 liver, 15 stomach, and 18 colon pairs) was utilized for tissue-level validation. Detailed information on all datasets and their sources are provided in **Supplementary Table 2**. Detailed clinical characteristics of the patients and healthy controls across these cohorts are summarized in **Supplementary Tables 13-19**. For the single-end WGBS dataset comprising LIHC and healthy cfDNA samples, raw reads were filtered using fastp (v0.22.0; *-f 8 -q 20*)^61^ to retain high-quality reads. For all other pair-end datasets, raw reads were trimmed and filtered using Trim Galore (v0.6.1)^62^ with parameter settings consistent with the original studies. Then, trimmed reads were aligned to the human reference genome (hg38) using Bismark (v0.24.2)^63^. For the EM-seq dataset, only Bismark-aligned BAM files mapped to hg19 were available, which were converted to hg38 coordinates using CrossMap (v0.6.6)^64^ to ensure genome build consistency. The aligned reads were subsequently filtered with Samtools (v1.6; *-q 10 - f 2 -F 1796*)^65^ to retain high-quality, properly mapped reads. For cfMethyl-Seq and RRBS tissue datasets containing Unique Molecular Identifier (UMI), PCR duplicates were removed using Umi-Grinder (v0.2.0)^66^. For all remaining datasets, PCR duplicates were removed using the *deduplicate_bismark* function in Bismark (v0.24.2)^63^. CpG methylation calls were then extracted using the *bismark_methylation_extractor* function in Bismark (v0.24.2)^63^. The resulting methylation call files were used as inputs for downstream cfMIND analyses.

### Assessment of cfMIND under different sequencing depth

We selected the largest cohort from the cfMethyl-Seq dataset, comprising 67 LUAD and 193 healthy samples. BAM files were down-sampled to simulate sequencing depths ranging from 0.2× to 20× using Samtools (v1.6)^65^ with random reads sampling. Down-sampling at each depth was independently repeated five times using different random seeds. Each dataset, corresponding to a specific sequencing depth and random seed, was then analyzed using the standard cfMIND pipeline. Model performance was evaluated using leave-one-out cross-validation. The AUROC was calculated for each run, and the average AUROC across the five repeats was recorded for each sequencing depth.

### Processing workflow for comparator methods

- Baseline_DMR was implemented by performing DMR calling with Metilene (v0.2-8)^67^ on the same genomic bins utilized by cfMIND, applying the following criteria: a q-value threshold of 0.05, a minimum of three CpGs per region, and a minimum methylation difference of 0.1 between groups. The resulting DMRs were subjected to the same feature selection and model construction procedures as cfMIND.
- CancerDetector was executed following its guideline. LIHC detection in Fig. 2 was performed using the liver-cancer-specific markers from the original study^16^. When applied to other cancer types, markers were identified using methylation sequencing data from tumor and normal tissue samples (**Supplementary Table 2**: EGAS00001006020) together with healthy plasma samples (**Supplementary Table 2**: EGAS00001000566), using a custom script that implemented the marker identification strategy described in the original CancerDetector study. All subsequent analyses were performed using the cfTools R package (v1.10.0)^68^, including input data preparation, retention of cfDNA fragments overlapping the markers, and application of CancerDetector to estimate tumor fraction.
- DISMIR was executed following its guideline. LIHC detection in Fig. 2 was performed using the liver-cancer-specific markers from the original study^17^. When applied to other cancer types, cancer-specific DMRs were first identified with methylation sequencing data from tumor tissue samples (**Supplementary Table 2**: EGAS00001006020) and healthy plasma samples (**Supplementary Table 2**: EGAS00001000566) using the *get_switching_region* function provided by DISMIR. Plasma cfDNA reads overlapping these DMRs were then selected using the *generate_new_bam* and *extract_read_from_bam* functions. Subsequently, a deep learning model was trained using the *DISMIR_training* function. Finally, DISMIR_predict_reads_source and DISMIR_cal_risk scripts were used to estimate tumor fraction.

### Integrated evaluation of computational efficiency and classification performance

The computational efficiency of cfMIND and comparator methods was evaluated using the targetBS dataset by recording runtime, peak CPU memory usage, and peak GPU memory usage using the Slurm *sacct* command. All methods were executed on a Linux (CentOS 7)-based high-performance computing node equipped with three NVIDIA A30 GPUs, 128 CPU cores, and ∼257 GB of system memory, with one GPU, 100 CPU threads, and 200 GB of memory allocated per run to ensure reliable evaluation. In parallel, classification performance was assessed using the mean AUROC for each method, calculated based on the datasets shown in **Fig. 2**. To enable fair integration of performance and computational efficiency, all metrics were transformed using min-max normalization and rescaled to values between 0 and 1. For metrics in which smaller values indicate better computational efficiency (runtime, CPU memory usage, and GPU memory usage), normalized scores were computed as: 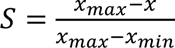. For metrics where larger values indicate better performance (AUROC), normalized scores were computed as: 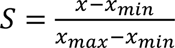, where *x* denotes the metric value, and *x*_*max*_ and *x*_*min*_ represent the minimum and maximum values of that metric across all methods. Based on all normalized metrics, the composite score for a method was defined as: *Composite Score*; 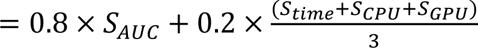.

### Gene mapping and functional enrichment analysis

Genomic annotation of cfMIND feature regions was performed using the ChIPseeker R package (v1.38.0; *stssRegion=c(-3000, 3000) TxDb=txdb annoDb=“org.Hs.eg.db”*). Gene Ontology (GO) enrichment analysis was conducted using the *enrichGO* function in the clusterProfiler R package (v4.10.1; *OrgDb=org.Hs.eg.db ont=“BP” readable=TRUE pAdjustMethod=“BH” pvalueCutoff=0.05*). Gene Set Enrichment Analysis (GSEA) was performed using the *GSEA* function in the clusterProfiler R package (v4.10.1; *pvalueCutoff=0.05 scoreType=“std”*), with genes ranked in descending order based on the *log*_2_(*fold change*) of feature values between the compared groups. Enriched gene sets with adjusted p values < 0.05 were deemed significant.

### Tissue and single cell level analysis of cfMIND features in COAD

In cfMIND-based COAD detection, the top 5,000 features were selected based on importance, and features exhibiting significant *M*_0′ and *M*_1′ differences between COAD and normal cfDNA samples were retained. These features were then categorized into four groups by comparing the mean feature values between COAD and normal samples: *M_0.TvsN.down* (lower *M*_0′ in tumors), *M_0.TvsN.up* (higher *M*_0′ in tumors), *M_1.TvsN.down* (lower *M*_1′ in tumors), and *M_1.TvsN.up* (higher *M*_1′ in tumors), with each group corresponding to a distinct set of genomic regions.

### · TCGA tissue DNA methylation array and RNA-seq data analysis

Tissue DNA methylation array data (n=349, comprising 311 tumor and 38 normal tissues) from the TCGA COAD cohort (**Supplementary Table 2**) were used for the analysis in this study. cfMIND feature regions containing fewer than three CpGs were excluded. Average methylation levels for the remaining regions were calculated and compared between tumor and normal tissues. Expression data (TPM) based on bulk RNA-seq (n=524, comprising 483 tumor and 41 normal tissues) from TCGA COAD cohort were used for the analysis in this study (**Supplementary Table 2**). For genes whose promoters mapped with the feature regions used in above tissue DNA methylation analysis, expression levels were compared using TPM values standardized to z-scores within each sample.

### · Single-cell bisulfite sequencing data analysis

Processed single-cell bisulfite sequencing data were downloaded from GSE97693^42^. Patient CRC01 with the largest sequenced cell numbers was used, spanning COAD sublineages A0-A9 and B0-B1 as defined in the original study. For each cell, the mean methylation level for each region in the four feature groups was computed. Each COAD sublineage was then compared with the normal colon (NC) group for regions that contained more than two non-missing values in both groups using a two-sided Wilcoxon rank-sum test. Focusing on feature groups with more than 100 regions (*M_0.TvsN.down* and *M_1.TvsN.up*), we selected the top 2 comparisons with significantly more upregulated features than downregulated features compared with NC. The resulted −*log*_10_(*p value*) and *log*_2_*FC* for all retained regions were visualized using volcano plots.

### · Single-cell RNA-seq data analysis

Single-cell UMI count matrices for COAD samples were downloaded from CancerSCEM (https://ngdc.cncb.ac.cn/cancerscem/) database with per-cell type annotations. Data from 34 tumor samples and 23 adjacent normal tissue samples with more than 700 cells were used for the downstream analysis^43–47^. The count matrix was processed using the Seurat R package (v5)^69^. Cells with <200 or >5,000 detected genes or ≥10% mitochondrial gene expression were removed. Doublets were identified and excluded with DoubletFinder^70^ (v2.0.6; expected_doublet_rate = 0.075 PCs=1:10 pN=0.25 pK= optimal_pk resolution=0.5). Cells were annotated with the original cell types provided by CancerSCEM. Meta-gene expression for each feature group was calculated per cell using the *AddModuleScore* function in Seurat.

## Author contributions

Y.L. conceived and supervised this project. J.L. and Z.L. performed most of the analysis. Z. H. and Y. Z. performed part of the analysis. Y.L. wrote most part of the manuscript. All authors interpreted the results, read, revised, and approved the final manuscript.

## Acknowledgments

The authors thank members of Yumei Li’s lab and Wei Li’s lab for helpful discussions.

## Supplementary information

Supplementary Figs. 1 to 9, and Tables 1 to 19.

## References

1 Ma, L. et al. Liquid biopsy in cancer current: status, challenges and future prospects. Signal Transduct Target Ther 9, 336 (2024). 10.1038/s41392-024-02021-w

2 Cohen, J. D. et al. Detection and localization of surgically resectable cancers with a multi-analyte blood test. Science 359, 926–930 (2018). 10.1126/science.aar3247

3 Cristiano, S. et al. Genome-wide cell-free DNA fragmentation in patients with cancer. Nature 570, 385–389 (2019). 10.1038/s41586-019-1272-6

4 Foda, Z. H. et al. Detecting Liver Cancer Using Cell-Free DNA Fragmentomes. Cancer Discov 13, 616–631 (2023). 10.1158/2159-8290.Cd-22-0659

5 Conway, A. M. et al. A cfDNA methylation-based tissue-of-origin classifier for cancers of unknown primary. Nat Commun 15, 3292 (2024). 10.1038/s41467-024-47195-7

6 Sadeh, R. et al. ChIP-seq of plasma cell-free nucleosomes identifies gene expression programs of the cells of origin. Nat Biotechnol 39, 586–598 (2021). 10.1038/s41587-020-00775-6

7 Huang, J. & Wang, L. Cell-Free DNA Methylation Profiling Analysis-Technologies and Bioinformatics. Cancers (Basel) 11 (2019). 10.3390/cancers11111741

8 Pietrzak, B., Kawacka, I., Olejnik-Schmidt, A. & Schmidt, M. Circulating Microbial Cell-Free DNA in Health and Disease. Int J Mol Sci 24 (2023). 10.3390/ijms24033051

9 Stejskal, P. et al. Circulating tumor nucleic acids: biology, release mechanisms, and clinical relevance. Mol Cancer 22, 15 (2023). 10.1186/s12943-022-01710-w

10 Moss, J. et al. Comprehensive human cell-type methylation atlas reveals origins of circulating cell-free DNA in health and disease. Nat Commun 9, 5068 (2018). 10.1038/s41467-018-07466-6

11 Nassiri, F. et al. Detection and discrimination of intracranial tumors using plasma cell-free DNA methylomes. Nat Med 26, 1044–1047 (2020). 10.1038/s41591-020-0932-2

12 Jamshidi, A. et al. Evaluation of cell-free DNA approaches for multi-cancer early detection. Cancer Cell 40, 1537–1549.e1512 (2022). 10.1016/j.ccell.2022.10.022

13 Widschwendter, M. et al. The potential of circulating tumor DNA methylation analysis for the early detection and management of ovarian cancer. Genome Med 9, 116 (2017). 10.1186/s13073-017-0500-7

14 Chen, X. et al. Non-invasive early detection of cancer four years before conventional diagnosis using a blood test. Nat Commun 11, 3475 (2020). 10.1038/s41467-020-17316-z

15 Newman, A. M. et al. An ultrasensitive method for quantitating circulating tumor DNA with broad patient coverage. Nat Med 20, 548–554 (2014). 10.1038/nm.3519

16 Li, W. et al. CancerDetector: ultrasensitive and non-invasive cancer detection at the resolution of individual reads using cell-free DNA methylation sequencing data. Nucleic Acids Res 46, e89 (2018). 10.1093/nar/gky423

17 Li, J. et al. DISMIR: Deep learning-based noninvasive cancer detection by integrating DNA sequence and methylation information of individual cell-free DNA reads. Brief Bioinform 22 (2021). 10.1093/bib/bbab250

18 Jeong, Y. et al. MethylBERT enables read-level DNA methylation pattern identification and tumour deconvolution using a Transformer-based model. Nat Commun 16, 788 (2025). 10.1038/s41467-025-55920-z

19 Lister, R. et al. Human DNA methylomes at base resolution show widespread epigenomic differences. Nature 462, 315–322 (2009). 10.1038/nature08514

20 Cokus, S. J. et al. Shotgun bisulphite sequencing of the Arabidopsis genome reveals DNA methylation patterning. Nature 452, 215–219 (2008). 10.1038/nature06745

21 Stackpole, M. L. et al. Cost-effective methylome sequencing of cell-free DNA for accurately detecting and locating cancer. Nat Commun 13, 5566 (2022). 10.1038/s41467-022-32995-6

22 Vaisvila, R. et al. Enzymatic methyl sequencing detects DNA methylation at single-base resolution from picograms of DNA. Genome Res 31, 1280–1289 (2021). 10.1101/gr.266551.120

23 Siejka-Zielińska, P. et al. Cell-free DNA TAPS provides multimodal information for early cancer detection. Sci Adv 7, eabh0534 (2021). 10.1126/sciadv.abh0534

24 Bartolomucci, A. et al. Circulating tumor DNA to monitor treatment response in solid tumors and advance precision oncology. NPJ Precis Oncol 9, 84 (2025). 10.1038/s41698-025-00876-y

25 Swanton, C. Intratumor heterogeneity: evolution through space and time. Cancer Res 72, 4875–4882 (2012). 10.1158/0008-5472.Can-12-2217

26 Zhang, J. et al. Intratumor heterogeneity in localized lung adenocarcinomas delineated by multiregion sequencing. Science 346, 256–259 (2014). 10.1126/science.1256930

27 Anusewicz, D., Orzechowska, M. & Bednarek, A. K. Lung squamous cell carcinoma and lung adenocarcinoma differential gene expression regulation through pathways of Notch, Hedgehog, Wnt, and ErbB signalling. Sci Rep 10, 21128 (2020). 10.1038/s41598-020-77284-8

28 Asamura, H. et al. A Japanese Lung Cancer Registry study: prognosis of 13,010 resected lung cancers. J Thorac Oncol 3, 46–52 (2008). 10.1097/JTO.0b013e31815e8577

29 Herbst, R. S., Morgensztern, D. & Boshoff, C. The biology and management of non-small cell lung cancer. Nature 553, 446–454 (2018). 10.1038/nature25183

30 Jin, Y. et al. Whole-genome bisulfite sequencing of cell-free DNA unveils age-dependent and ALS-associated methylation alterations. Cell Biosci 15, 26 (2025). 10.1186/s13578-025-01366-1

31 Ilieva, H., Vullaganti, M. & Kwan, J. Advances in molecular pathology, diagnosis, and treatment of amyotrophic lateral sclerosis. Bmj 383, e075037 (2023). 10.1136/bmj-2023-075037

32 Shimizu, M. et al. RGMa collapses the neuronal actin barrier against disease-implicated protein and exacerbates ALS. Sci Adv 9, eadg3193 (2023). 10.1126/sciadv.adg3193

33 Sung, H. et al. Global Cancer Statistics 2020: GLOBOCAN Estimates of Incidence and Mortality Worldwide for 36 Cancers in 185 Countries. CA Cancer J Clin 71, 209–249 (2021). 10.3322/caac.21660

34 Leiphrakpam, P. D. & Are, C. PI3K/Akt/mTOR Signaling Pathway as a Target for Colorectal Cancer Treatment. Int J Mol Sci 25 (2024). 10.3390/ijms25063178

35 Sanaei, M. J. et al. The PI3K/Akt/mTOR axis in colorectal cancer: Oncogenic alterations, non-coding RNAs, therapeutic opportunities, and the emerging role of nanoparticles. J Cell Physiol 237, 1720–1752 (2022). 10.1002/jcp.30655

36 Narayanankutty, A. PI3K/ Akt/ mTOR Pathway as a Therapeutic Target for Colorectal Cancer: A Review of Preclinical and Clinical Evidence. Curr Drug Targets 20, 1217–1226 (2019). 10.2174/1389450120666190618123846

37 Zhang, Z. et al. Upregulated miR-1258 regulates cell cycle and inhibits cell proliferation by directly targeting E2F8 in CRC. Cell Prolif 51, e12505 (2018). 10.1111/cpr.12505

38 Malumbres, M. & Barbacid, M. Cell cycle, CDKs and cancer: a changing paradigm. Nat Rev Cancer 9, 153–166 (2009). 10.1038/nrc2602

39 Weinstein, J. N. et al. The Cancer Genome Atlas Pan-Cancer analysis project. Nat Genet 45, 1113–1120 (2013). 10.1038/ng.2764

40 Newell-Price, J., Clark, A. J. & King, P. DNA methylation and silencing of gene expression. Trends Endocrinol Metab 11, 142–148 (2000). 10.1016/s1043-2760(00)00248-4

41 Heery, R. & Schaefer, M. H. Systematic identification of regions where DNA methylation is correlated with transcription refines regulatory logic in normal and tumour tissues. Nucleic Acids Res 53 (2025). 10.1093/nar/gkaf949

42 Bian, S. et al. Single-cell multiomics sequencing and analyses of human colorectal cancer. Science 362, 1060–1063 (2018). 10.1126/science.aao3791

43 Lee, H. O. et al. Lineage-dependent gene expression programs influence the immune landscape of colorectal cancer. Nat Genet 52, 594–603 (2020). 10.1038/s41588-020-0636-z

44 Qian, J. et al. A pan-cancer blueprint of the heterogeneous tumor microenvironment revealed by single-cell profiling. Cell Res 30, 745–762 (2020). 10.1038/s41422-020-0355-0

45 Becker, W. R. et al. Single-cell analyses define a continuum of cell state and composition changes in the malignant transformation of polyps to colorectal cancer. Nat Genet 54, 985–995 (2022). 10.1038/s41588-022-01088-x

46 Bala, P. et al. Aberrant cell state plasticity mediated by developmental reprogramming precedes colorectal cancer initiation. Sci Adv 9, eadf0927 (2023). 10.1126/sciadv.adf0927

47 Yang, M. et al. Single-cell analysis reveals cellular reprogramming in advanced colon cancer following FOLFOX-bevacizumab treatment. Front Oncol 13, 1219642 (2023). 10.3389/fonc.2023.1219642

48 Zhang, E. et al. Roles and mechanisms of tumour-infiltrating B cells in human cancer: a new force in immunotherapy. Biomark Res 11, 28 (2023). 10.1186/s40364-023-00460-1

49 Edin, S. et al. The Prognostic Importance of CD20(+) B lymphocytes in Colorectal Cancer and the Relation to Other Immune Cell subsets. Sci Rep 9, 19997 (2019). 10.1038/s41598-019-56441-8

50 Berntsson, J. et al. Expression of programmed cell death protein 1 (PD-1) and its ligand PD-L1 in colorectal cancer: Relationship with sidedness and prognosis. Oncoimmunology 7, e1465165 (2018). 10.1080/2162402x.2018.1465165

51 Chan, K. C. et al. Noninvasive detection of cancer-associated genome-wide hypomethylation and copy number aberrations by plasma DNA bisulfite sequencing. Proc Natl Acad Sci U S A 110, 18761–18768 (2013). 10.1073/pnas.1313995110

52 Lehmann-Werman, R. et al. Identification of tissue-specific cell death using methylation patterns of circulating DNA. Proc Natl Acad Sci U S A 113, E1826–1834 (2016). 10.1073/pnas.1519286113

53 Thierry, A. R., El Messaoudi, S., Gahan, P. B., Anker, P. & Stroun, M. Origins, structures, and functions of circulating DNA in oncology. Cancer Metastasis Rev 35, 347–376 (2016). 10.1007/s10555-016-9629-x

54 Schreiber, R. D., Old, L. J. & Smyth, M. J. Cancer immunoediting: integrating immunity’s roles in cancer suppression and promotion. Science 331, 1565–1570 (2011). 10.1126/science.1203486

55 Stanton, S. E. et al. SITC strategic vision: prevention, premalignant immunity, host and environmental factors. J Immunother Cancer 13 (2025). 10.1136/jitc-2024-010419

56 Jin, Y., Allen, E. G. & Jin, P. Cell-free DNA methylation as a potential biomarker in brain disorders. Epigenomics 14, 369–374 (2022). 10.2217/epi-2021-0416

57 Kim, K. et al. Cell-free DNA methylation-based inflammation score as a marker for hepatocellular carcinoma among people living with HIV. Hepatol Int 19, 596–606 (2025). 10.1007/s12072-024-10768-1

58 Mondelo-Macía, P., Castro-Santos, P., Castillo-García, A., Muinelo-Romay, L. & Diaz-Peña, R. Circulating Free DNA and Its Emerging Role in Autoimmune Diseases. J Pers Med 11 (2021). 10.3390/jpm11020151

59 Sun, K. et al. Orientation-aware plasma cell-free DNA fragmentation analysis in open chromatin regions informs tissue of origin. Genome Res 29, 418–427 (2019). 10.1101/gr.242719.118

60 Doebley, A. L. et al. A framework for clinical cancer subtyping from nucleosome profiling of cell-free DNA. Nat Commun 13, 7475 (2022). 10.1038/s41467-022-35076-w

61 Chen, S., Zhou, Y., Chen, Y. & Gu, J. fastp: an ultra-fast all-in-one FASTQ preprocessor. Bioinformatics 34, i884–i890 (2018). 10.1093/bioinformatics/bty560

62 Krueger, F. Trim Galore. (2021). https://www.bioinformatics.babraham.ac.uk/projects/trim_galore/

63 Krueger, F. & Andrews, S. R. Bismark: a flexible aligner and methylation caller for Bisulfite-Seq applications. Bioinformatics 27, 1571–1572 (2011). 10.1093/bioinformatics/btr167

64 Zhao, H. et al. CrossMap: a versatile tool for coordinate conversion between genome assemblies. Bioinformatics 30, 1006–1007 (2014). 10.1093/bioinformatics/btt730

65 Li, H. et al. The Sequence Alignment/Map format and SAMtools. Bioinformatics 25, 2078–2079 (2009). 10.1093/bioinformatics/btp352

66 Krueger, F. Unique Molecule Identifiers (UMIs) based sequencing deduplication software. https://github.com/FelixKrueger/Umi-Grinder

67 Jühling, F. et al. metilene: fast and sensitive calling of differentially methylated regions from bisulfite sequencing data. Genome Res 26, 256–262 (2016). 10.1101/gr.196394.115

68 Ran, H. cfTools. (2025). https://bioconductor.org/packages/release/bioc/vignettes/cfTools/inst/doc/cfTools-vignette.html

69 Hao, Y. H. et al. Dictionary learning for integrative, multimodal and scalable single-cell analysis. Nature Biotechnology 42 (2024). 10.1038/s41587-023-01767-y

70 Stoeckius, M. et al. Cell Hashing with barcoded antibodies enables multiplexing and doublet detection for single cell genomics. Genome Biology 19 (2018). 10.1186/s13059-018-1603-1

