## Supplementary figures for "cfMIND: A read-level methylation framework for accurate non-invasive disease detection using cell-free DNA"

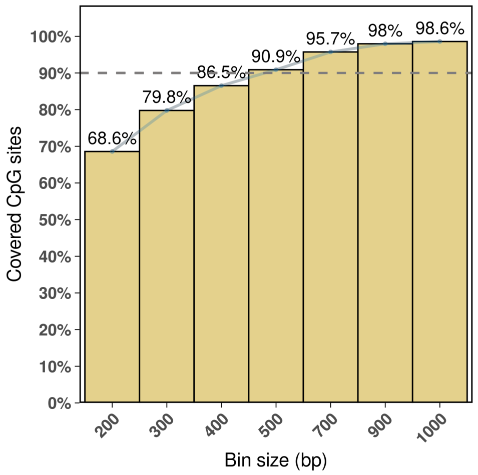


**Supplementary Fig. 1. Proportion of CpG sites retained under different genomic bin sizes.** The genome was binned into regions of 200-1000 bp, and bins containing ≥ 3 CpG sites were retained. The figure shows the fraction of CpG sites preserved after filtering, which exceeds 90% when the bin size is greater than 500 bp.

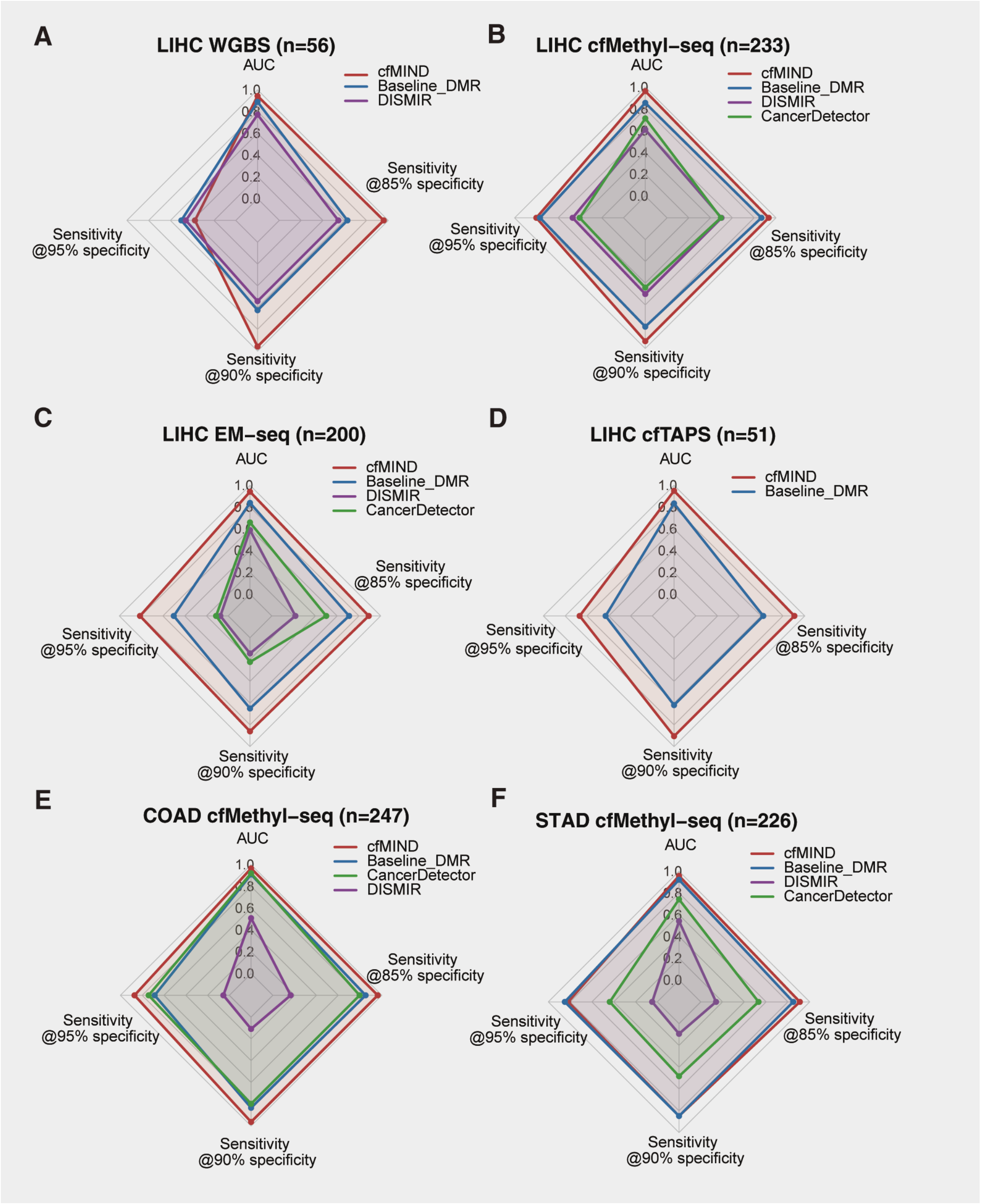


**Supplementary Fig. 2. Performance of cfMIND in cancer detection across multiple sequencing technologies and cancer types.**

**(A-D)** Radar charts presenting AUC and Sensitivity at 95%, 90%, and 85% specificity for LIHC detection using cfMIND, Baseline_DMR, CancerDetector, and DISMIR across four cfDNA methylation sequencing technologies: WGBS **(A)**, cfMethyl-Seq **(B)**, EM-seq **(C)**, and cfTAPS **(D)**.

**(E-F)** Radar charts presenting AUC and Sensitivity at 95%, 90%, and 85% specificity for COAD **(E)** and STAD **(F)** detection using cfMIND and comparator methods. All results were obtained using leave-one-out cross-validation, and the number of samples for each dataset was indicated in parentheses.

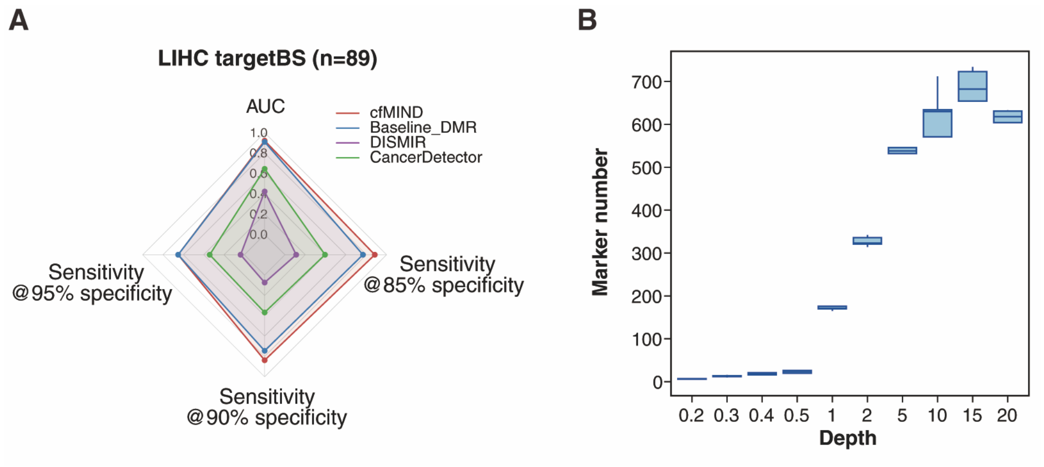


**Supplementary Fig. 3. Performance of cfMIND for LIHC detection in a targeted bisulfite sequencing dataset (targetBS) and at varying sequencing depths.**

**(A)** Radar chart presenting AUC and Sensitivity at 95%, 90%, and 85% specificity for detecting LIHC in a targeted bisulfite sequencing dataset.

**(B)** Boxplots showing feature numbers of LUAD detection at varying sequencing depths, assessed by LOOCV with five replicates per depth via subsampling.

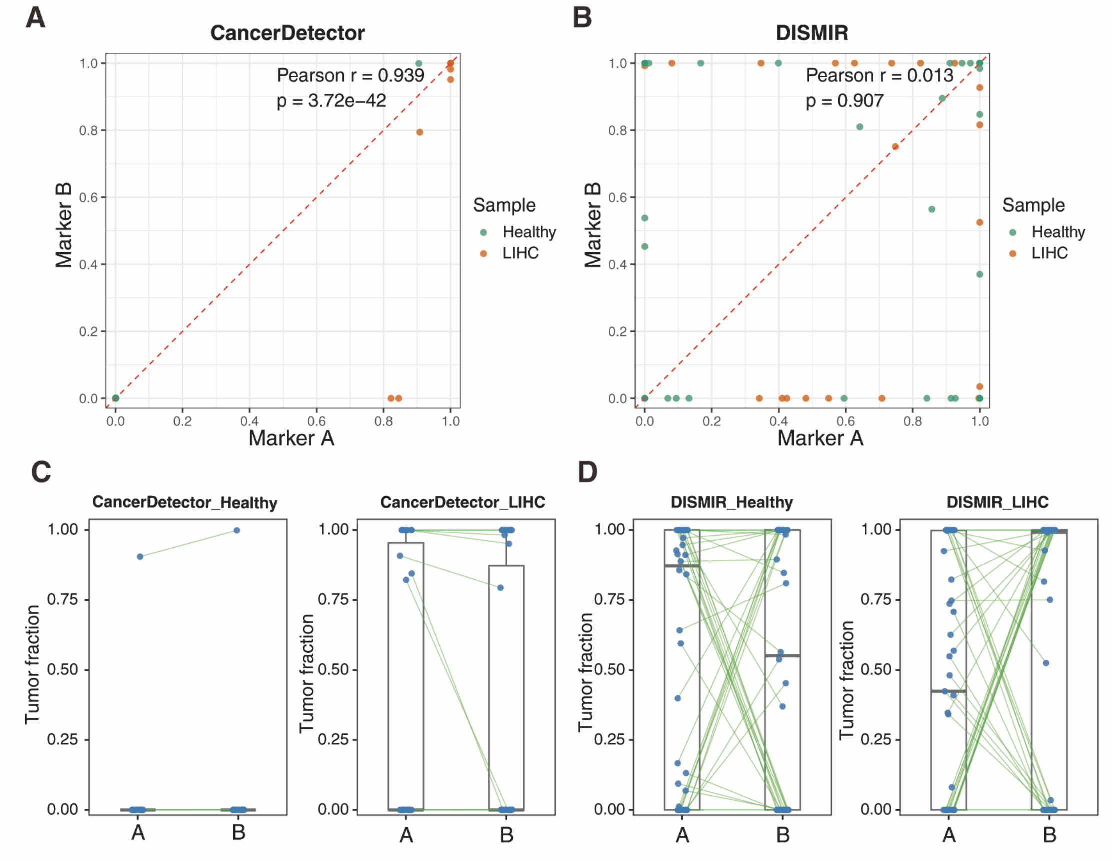


**Supplementary Fig. 4. Performance of CancerDetector and DISMIR across different tumor tissue datasets.**

**(A-B)** Correlation of prediction results on the targetBS dataset using marker set A and marker set B identified from different tissue datasets by CancerDetector **(A)** and DISMIR **(B)**. Each point represents a sample (healthy in green, LIHC in orange), with the Pearson correlation coefficient (r) and corresponding p-value indicated.

**(C-D)** Connected boxplots illustrating the prediction results of CancerDetector **(C)** and DISMIR **(D)** in predicting healthy and LIHC samples using two distinct marker sets (A and B) within the targetBS dataset.

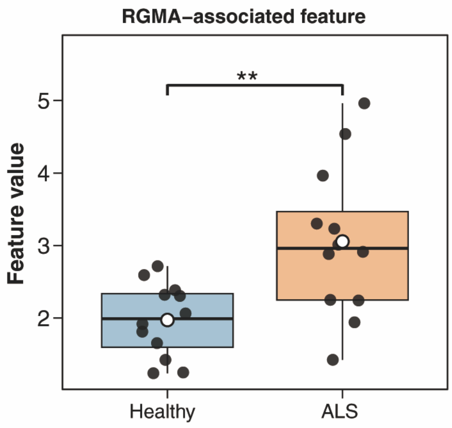


**Supplementary Fig. 5.** Boxplots showing the feature values for the RGMA-associated feature in ALS patients and healthy controls. Statistical significance was assessed using a two-sided Wilcoxon rank-sum test.

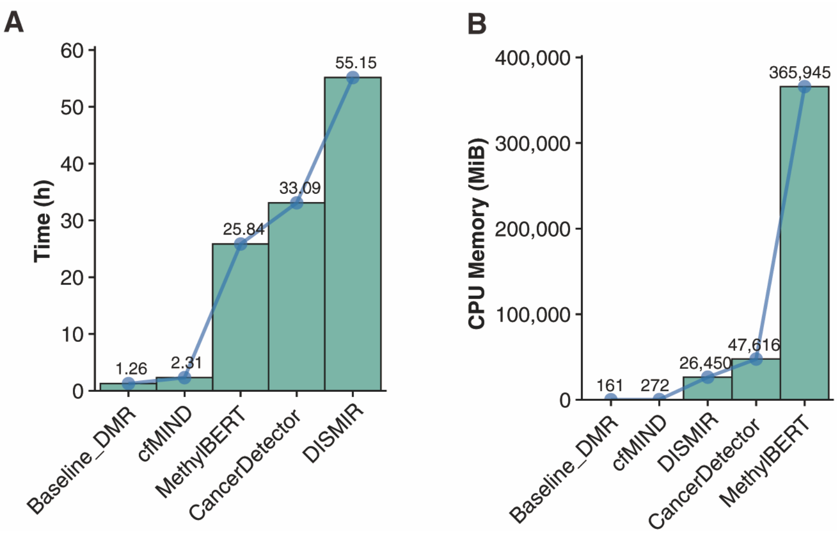


**Supplementary Fig. 6.** Comparison of computational efficiency between cfMIND and other methods, with runtime (hours) for leave-one-out cross-validation **(A)** and peak CPU memory usage (MiB) during each execution **(B)**.

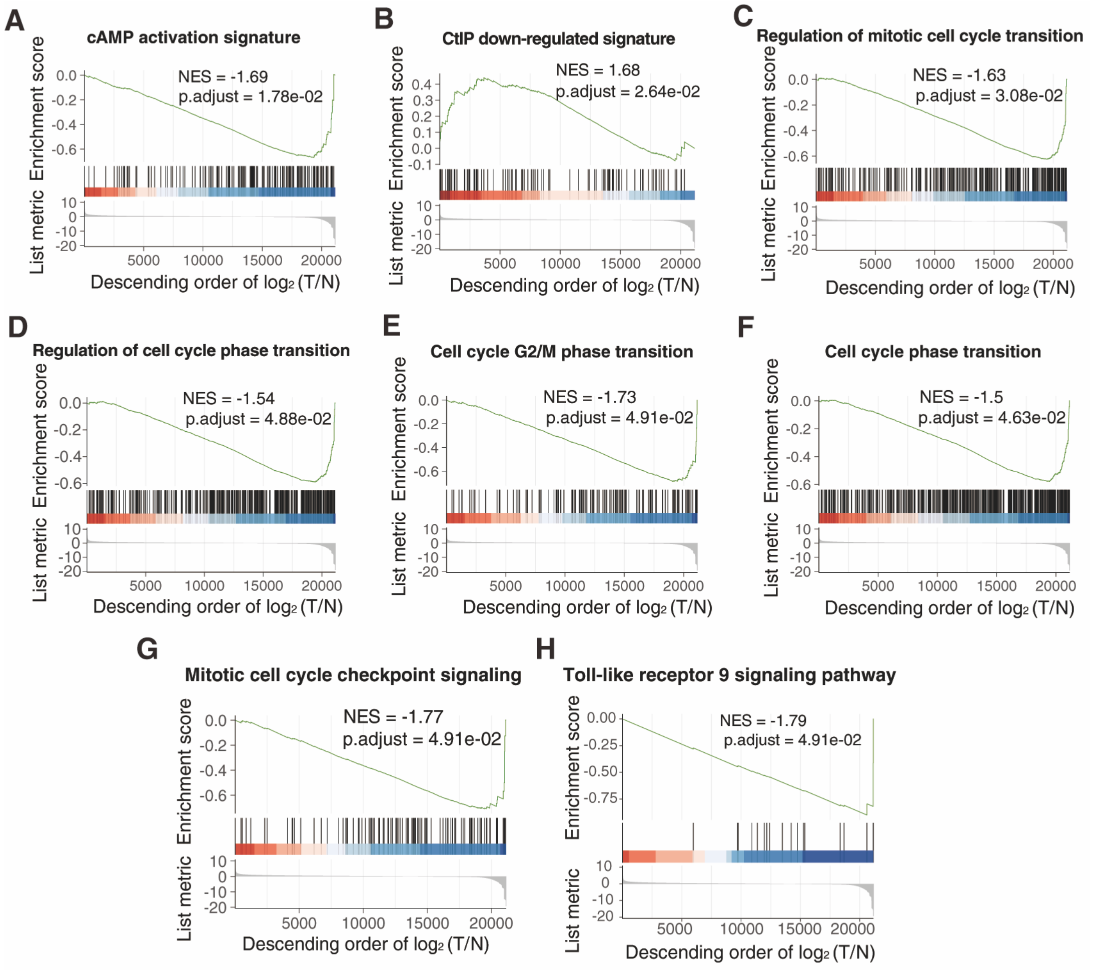


**Supplementary Fig. 7.** Enrichment plots showing the cancer-relevant GSEA results of cfMIND-derived features based on MSigDB C6 oncogenic signatures (A-B) and MSigDB C5 GO biological process pathways (C-H).

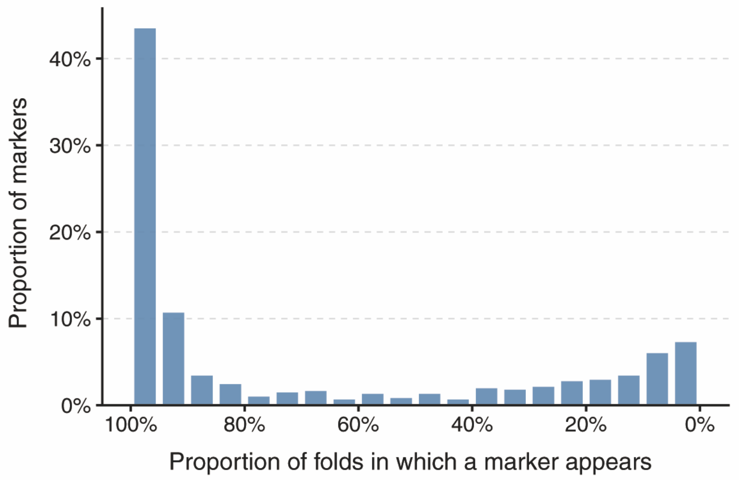


**Supplementary Fig. 8. Stability of COAD markers across leave-one-out cross-validation folds.** The bar plot shows the distribution of COAD markers according to their selection frequency across leave-one-out cross-validation (LOOCV) folds. The x-axis represents the proportion of LOOCV folds in which a marker was selected (binned at 5% intervals), and the y-axis indicates the proportion of markers within each bin.

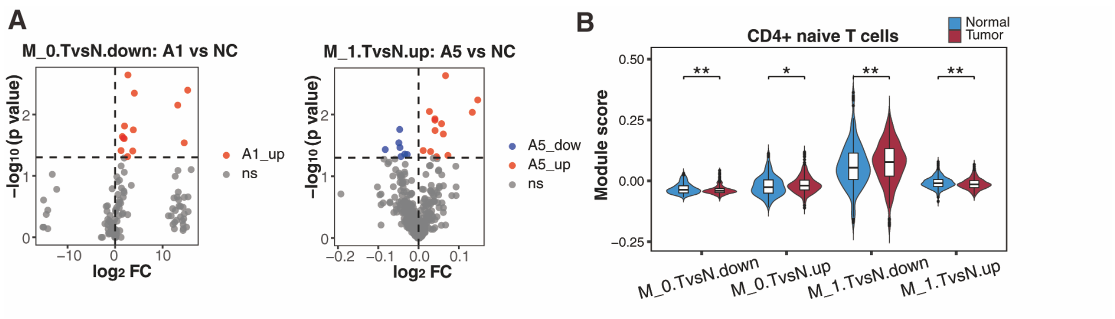


**Supplementary Fig. 9. Characterization of cfMIND-derived features at the single-cell level.**

**(A)** Volcano plots showing differentially methylated regions within the *M_0.TvsN.down* and *M_1.TvsN.up* sets at the single-cell level. Red dots indicate regions that are significantly hypermethylated in the corresponding COAD lineage, blue dots indicate significantly hypomethylated regions, and gray dots represent regions without significant methylation changes.

**(B)** Violin plots comparing gene expression levels between normal (blue) and tumor (red) $CD4^+$ naive T cells for genes mapped to promoter regions in the four sets: *M_0.TvsN.down*, *M_0.TvsN.up*, *M_1.TvsN.down*, and *M_1.TvsN.up*, with significance assessed by one-sided Wilcoxon rank-sum tests.
